# Genome size variation and evolution during invasive range expansion in an introduced plant

**DOI:** 10.1101/2022.08.16.504051

**Authors:** F. Alice Cang, Shana R. Welles, Jenny Wong, Maia Ziaee, Katrina M. Dlugosch

**Affiliations:** University of Arizona, Tucson, AZ, USA; Utah Valley University, Orem, UT, USA; Mills College, Oakland, CA, USA

## Abstract

Plants demonstrate some of the greatest variation in genome size among eukaryotes, and their genome sizes can vary dramatically across individuals and populations within species. This genetic variation can have consequences for traits and fitness, but few studies have been able to attribute genome size differentiation to ecological and evolutionary processes. Biological invasions present particularly useful natural laboratories to infer selective agents that might drive genome size shifts across environments and population histories. Here, we test hypotheses for the evolutionary causes of genome size variation across 14 invading populations of yellow starthistle, *Centaurea solstitialis*, in California, USA. We use a survey of genome sizes and trait variation to ask: (1) Is variation in genome size associated with developmental trait variation? (2) Are genome sizes smaller toward the leading edge of the expansion, consistent with selection for ‘colonizer’ traits? Or alternatively, does genome size increase toward the leading edge of the expansion, consistent with predicted consequences of founder effects and drift? (3) Finally, are genome sizes smaller at higher elevations, consistent with selection for shorter development times? We found that 2C DNA content varied 1.21-fold among all samples, and was associated with flowering time variation, such that plants with larger genomes reproduced later, with lower lifetime capitula production. Genome sizes increased toward the leading edge of the invasion, but tended to decrease at higher elevations, consistent with genetic drift during range expansion but potentially strong selection for smaller genomes and faster development time at higher elevations.

## Introduction

Plants demonstrate some of the greatest variation in genome size among eukaryotes (Bennetzen *et al*. 2005) and the potential ecological and evolutionary consequences of this widespread variation remain an open question. At the most basic level, we expect that large genomes require greater cell volumes and longer replication times, and comparative analyses across plants bear out the generality of this relationship, along with associated reductions in stomatal density and increased seed mass (Beaulieu *et al*. 2007a; Beaulieu *et al*. 2008). These differences suggest potential costs associated with maintaining large genomes, as greater cellular volumes and longer cell cycles might constrain functional traits related to growth, reproduction and dispersal, including longer minimum generation times (Grotkopp *et al*. 2004), slower rates of photosynthesis (Knight *et al*. 2005) and smaller plant size (Carta and Peruzzi 2016), among others (e.g. Kenton *et al*. 1986; Francis *et al*. 2008; Vesely *et al*. 2020). While there has been some success in identifying these effects at the cellular level and subsequent trait changes (Knight and Beaulieu 2008; Hodgson *et al*. 2010), population consequences and fitness effects in particular environments remain less understood (Gaut and Ross-Ibarra 2008; Whitney *et al*. 2010; Mei *et al*. 2017; Bilinski *et al*. 2018).

Contrary to traditional understanding of genome size as a stable species-level trait, genome structure can vary dramatically across individuals and populations and over time, even on very short time scales. The predominant molecular mechanisms of genome expansion are whole genome or whole chromosome duplications, and transposable element (TE) proliferation within a ploidy level (Feschotte and Pritham 2007; Tenaillon *et al*. 2010; Lisch 2013). Conversely, unequal homologous recombination or illegitimate recombination drive losses of DNA content (Bennetzen *et al*. 2005). Some of the clearest evidence for the mutability of genome content and the processes that influence genome has been observed in crop systems and wild crop relatives. Some of the earliest evidence emerged from studies of wild barley, in which variation in insertion patterns of a single family, BARE-1, has contributed up to ∼5% of genome size variation (Kalendar *et al*. 2000). In maize, the most common TE families that make up nearly ∼70% of the genome have average insertion dates of less than 1 Mya, with some inserting as recently as 560 kya (Baucom *et al*. 2009). These recent bursts of TE proliferation are responsible for wide-ranging variation among maize individuals, and TE activity explains the observation of >20% difference in genome size between B73 individuals (Vielle-Calzada *et al*. 2009). Similar findings in flax have identified a comparable scale of variation, as well as inducible changes to genome size within one generation (Cullis 2005), suggesting relatively few meiotic events are required to generate dramatic variation among closely related individuals. In cultivated rice, recent transposition of LTR-RT transposable element families are responsible for genome size doubling, on the same scale of variation as whole genome duplications (Piegu *et al*. 2006). Work in select model systems has shown similar findings, including in *Arabidopsis* (Schmuths *et al*. 2004; Davison *et al*. 2007) and *Drosophila* species (Petrov 2003; Ellis *et al*. 2014).

Few studies have been able to clearly attribute genome size variation to ecological and evolutionary processes. The *Arabidopsis* (Lockton *et al*. 2008; Lockton and Gaut 2010) and *Drosophila* genetic model systems (Gonzalez *et al*. 2008a; Gonzalez *et al*. 2008b) have leveraged abundant sequence data and have identified signatures of both selection and drift in families of TEs known to contribute to variation in genome size. Again, crop systems also provide evolutionary insights, as in the genus *Oryza*, where TE variation was identified as the driver of divergence between different species, indicating that mechanisms generating genome size variation can also have important macroevolutionary implications (Zhang and Gao 2017). In wild barley populations, BARE-1 retroelement content varied across elevational and aridity gradients, suggesting an association with environmental conditions (Kalendar *et al*. 2000). Multiple systems have identified potential selective effects of elevation on genome size, potentially due to differences in growing season and the fitness effects of development time, as in maize (Bilinski *et al*. 2018), *Corchorus olitorius* (Benor *et al*. 2011), *Lagenaria siceraria* (Achigan-Dako *et al*. 2008), as well as a small number of non-crop systems such as *Dactylis glomerata* (Creber *et al*. 1994), and *Arachis duranensis* (Temsch and Greilhuber 2001), though other results are conflicting (Lysak *et al*. 2000; Oney-birol and Tabur 2018; Savas *et al*. 2019). Taken together, these studies suggest that intraspecific genome size variation arises because the molecular mechanisms that influence genome size and subsequent structural variation are subject to the same evolutionary processes as other mutations. Subsequent accumulation of population-level differences can be ecologically relevant and function as important sources of variation upon which selection can act. Yet, there is still a dearth of knowledge about the fitness effects of genome size variation in natural populations, and in non-model systems.

Biological invasions present particularly useful natural laboratories to examine ecological consequences of intraspecific genome size variation and to infer selective agents that might drive genome size shifts. If genome size imposes developmental constraints, a potential invader may benefit from reduced genome sizes that promote fast generation time and high reproductive rates, which are traits associated with a colonizer life history (Baker 1974). Thus, genome size-associated traits that equip individuals to disperse, survive and reproduce in marginal habitats, may select for smaller genomes. This may account for the overrepresentation of species with small genomes among weedy taxa in broad-scale surveys of plant invaders globally (Kubesova *et al*. 2010; Pandit *et al*. 2014) and across the US (Kuester *et al*. 2014). A comparative phylogenetic study of pines identified relationships between small genomes and “weedy” traits, such as small seed size, short minimum generation time and fast relative growth rate, while also linking genome size to invasiveness as a character (Grotkopp *et al*. 2004). Importantly, invaded systems allow inter- and intra-population comparisons of introductions with their source populations, to infer contemporary and rapid evolution of genome size. Direct comparisons between introduced populations and known sources are relatively few in number (Crosby *et al*. 2014; Pysek *et al*. 2018) and methodological issues with genome size estimation can produce contested results (Lavergne *et al*. 2009; Martinez *et al*. 2018). Identifying evidence for selection on genome size during colonization requires systems in which the history of expansion is well-documented, ecologically relevant traits are known, and robust genome size estimation methods are used.

In contrast to natural selection, neutral population genetic processes might also influence genome size variation. If processes such as TE proliferation generate genetic variants that are neutral or only weakly deleterious, genetic drift may allow sequences that increase average genome size to persist and spread. This is especially important in invading populations, where range expansion can diminish the effective population size of successive founding events at the leading edge (Slatkin and Excoffier 2012; Braasch *et al*. 2019), and mutations arising in advancing populations can ‘surf’ to higher frequency and broader geographic scales (Klopfstein *et al*. 2005; Gralka *et al*. 2016), even if these variants are disadvantageous (Peischl *et al*. 2013). Thus, if drift and founder effects are stronger than selection, we expect genome sizes to increase as an invasion expands across space.

Here, we investigate evolutionary causes of genome size variation in the invasive species yellow starthistle, *Centaurea solstitialis* L., in the Asteraceae (hereafter, YST). YST is an annual, obligately outcrossing, diploid thistle (2*n*=16, Heiser and Whitaker 1948; Widmer *et al*. 2007; Öztürk *et al*. 2009), and previous work has suggested that genome size is variable within its invaded ranges (Irimia *et al*. 2017). YST is invasive in the United States, where it is particularly problematic in California and now occupies over 12 million acres of land (Maddox and Mayfield 1985; Pitcairn *et al*. 2006). YST likely originated in the eastern Mediterranean before an ancient expansion into Eurasia and western Europe (Barker *et al*. 2017). Historical records indicate that it was introduced to South America from Spain and to the US from Chile via imports of alfalfa, with the earliest recorded occurrence in the San Francisco Bay area in 1869 (Pitcairn *et al*. 2006). This recent colonization history has been confirmed by genetic studies (Dlugosch *et al*. 2013; Eriksen *et al*. 2014; Barker *et al*. 2017). We know from this previous work that Californian populations are derived from source populations in western Europe (Dlugosch *et al*. 2013; Barker *et al*. 2017) and have evolved shifts in growth rate and flowering time (Dlugosch *et al*. 2015), which are traits associated with genome size variation in other systems (eg. Grotkopp *et al*. 2004; Carta and Peruzzi 2015; Bilinski *et al* 2018). We also know that the invasion has crossed large elevational and environmental gradients (Gerlach 1997; Pitcairn *et al*. 2006; Braasch *et al*. 2019), which provide an opportunity to examine the influence of environmental clines on genome size variation, decoupled from invasion history. In particular, evidence from other plant systems (eg. Achigan-Dako *et al*. 2008; Benor *et al*. 2011; Bilinski *et al*. 2018) suggests higher elevations in particular may impose selection for smaller genome size

We leverage understanding of the expansion history of YST in California to test predictions about how genome size variation might have evolved under different evolutionary processes. If large genome size constrains growth and reproduction as observed in other systems, we expect selection to favor smaller genome sizes during the process of range expansion, and at high elevations regardless of expansion history. Alternatively, if neutral processes dominate, we expect genome sizes to increase during range expansion. We combine a survey of genome sizes estimated by flow cytometry with trait data collected from a common garden experiment to ask: (1) Is variation in genome size associated with developmental trait variation? (2) Are genome sizes smaller toward the leading edge of the expansion, consistent with selection for ‘colonizer’ traits? Or alternatively, does genome size increase toward the leading edge of the expansion, consistent with expectations of founder effects and drift? (3) Finally, are genome sizes associated with elevational clines that are decoupled from the direction of expansion?

## Materials and Methods

### Plant material

Seeds from introduced populations of YST from 14 sites in California (Fig. 1, Table 1) were collected in August 2016 along a linear transect at each site, with maternal plants >1m apart. Collections from California include populations from the coastal San Francisco Bay area which is the putative site of introduction to California, the Central Valley where the invasion is most severe and the Sierra Nevada mountain range, which includes the leading edge of the Californian expansion.

**Table 1.** Field collections from the YST invaded range in California, USA. Sample sizes of genome size estimates for each population include the overall number of individuals with at least 1 measurement. Values in parentheses indicate how many individuals out of the total have replicate estimates.

|  |  |  | Latitude, Longitude | Elevation (m) | N (Common garden) | N (Genome size) |
| --- | --- | --- | --- | --- | --- | --- |
| Coast | Diablo | DIA | N 37.863694, W 121.976444 | 237 | 20 | 21(1) |
|  | Gilroy | GIL | N 37.03373, W 121.53674 | 68 | 22 | 27(2) |
|  | Marin | MAR | N 38.01154, W 122.612222 | 143 | 25 | 26(2) |
|  | Napa | NAP | N 38.2400167, W 122.2702545 | 17 | 22 | 27(3) |
| Central Valley | Clovis | CLV | N 36.9231479, W 119.793704 | 118 | 17 | 18(2) |
|  | Red Bluff | RB | N 40.27083, W 122.27104 | 157 | 23 | 24(1) |
|  | Triangle | TRI | N 37.46178, W 119.79218 | 866 | 13 | 13 |
| Sierra Nevadas | Arnold | ARN | N 38.262671, W 120.336713 | 1273 | 30 | 30(2) |
|  | Belden | BEL | N 40.004779, W 121.261753 | 679 | 25 | 26(2) |
|  | Lyons | LYO | N 38.078308, W 120.164527 | 1497 | 21 | 21(1) |
|  | Placerville | PLA | N 38.733866, W 120.829686 | 507 | 21 | 23(2) |
|  | Quincy | QUI | N 39.964903, W 120.944909 | 1075 | 23 | 24(2) |
|  | Sandflat | SAN | N 38.76417, W 120.327177 | 1192 | 27 | 29(3) |
|  | Vet | VET | N 38.09996, W 120.58947 | 386 | 23 | 24(1) |
|  |  |  |  | Total | 312 | 333(24) |

**Figure 1.**
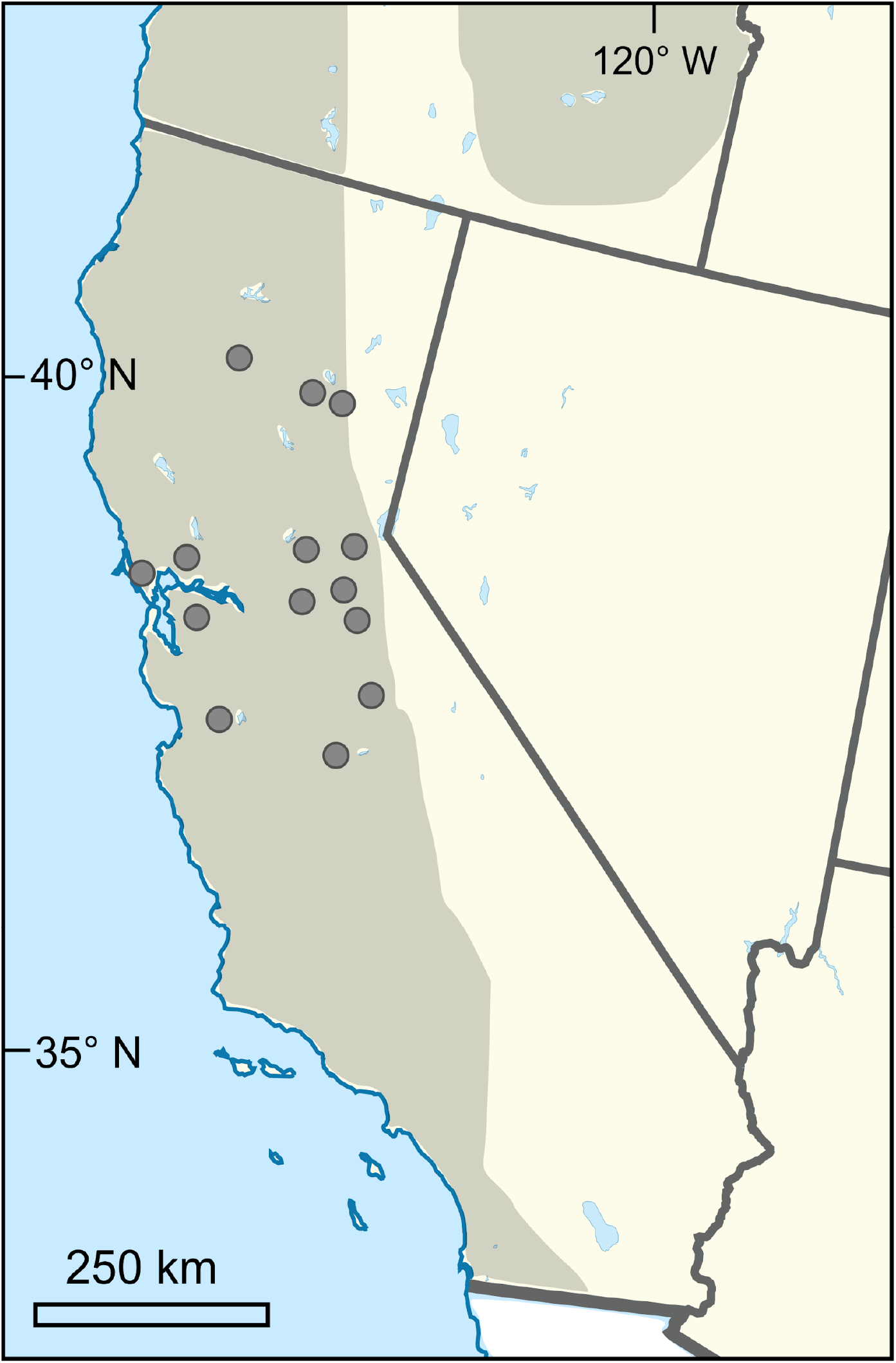
Map of 14 YST populations sampled in 2016 from the invasion in California, USA. Shaded areas indicate the invaded range in western North America.

### Common garden

We collected common garden data from 13-30 individuals from each of the 14 populations (N=312; Table 1). We germinated seeds on the surface of moist potting soil (3:2:1 ratio of Sunshine Mix #3 soil, vermiculite and 20 grit silica sand) under fluorescent lights and 12 hour days in December 2016 and recorded germination date daily. We germinated multiple seeds from the same maternal plant and kept the first germinating individual. We transplanted all growing plants into 410ml Deepots (Steuwe and Sons, Inc, USA) in January 2017 when they were 5 weeks old and grew them in a greenhouse at the University of Arizona in Tucson, AZ, USA. Once in the greenhouse, we randomly assigned one individual from each population to each of 30 different blocks. Additional individuals from two different maternal plants from ARN and SAN were transplanted into three and four blocks, respectively (Table 1). Plants were watered daily using an automatic drip watering system and maintained through senescence. At the rosette stage, prior to reproduction, we measured leaf number, length of longest leaf and width of longest leaf to estimate a size index and early growth rate (as in Dlugosch *et al*. 2015). The size index was calculated as (leaf number*(maximum leaf length*maximum leaf width)½), which correlates linearly with biomass in this species (Dlugosch *et al*. 2015). We calculated linear growth rates using our size index and number of days since germination. The common garden was checked daily to record days to the initiation of bolting and days until the first flower. Plants were harvested when most individuals had died, after which we counted the total number of flowering heads. We dried harvested plants overnight at 60°C and weighed aboveground dry biomass.

### Genome size estimation

We estimated genome size by flow cytometry using the FACSCanto II instrument (BD Biosciences, San Jose, CA, USA) equipped with a blue (488-nm), air-cooled, 20-mW solid state laser and a red (633-nm) 17-mW helium neon laser for UV excitation. Sample preparation followed a modified two-step protocol by Doležel *et al*. (2007) as follows. We chose *Raphanus sativus* (2C=1.11pg) provided by the Institute of Experimental Botany (Prague, Czech Republic) as our internal standard, as it has a close but non-overlapping genome size with the previously reported nuclear genome size of YST (2C=1.74pg; Bancheva and Greilhuber 2006). Nuclear suspensions were prepared by chopping 50-60 mg each of YST and *R. sativus* fresh tissue with 0.5 ml of ice-cold Otto I buffer in a Petri dish on top of ice, using a sharp razor blade. The suspension was filtered first through gauze and then again through 18 μm Nylon mesh. Nuclear suspensions in Otto I buffer are considered relatively stable (Doležel *et al*. 2007) and samples were left covered and on ice until immediately before analysis by flow cytometry, at which point we added 0.77ml of Otto II buffer, 29 μL of 1 mg/mL propidium iodide (PI) stain and 7.5 μL of 1 mg/mL RNase. The sample was then gently vortexed and left for at least 5 minutes on ice in a covered container. YST genome sizes were calculated according to Doležel and Bartos (2005). We recorded 3 estimates of genome size on different days for a subset of individuals (N=24), including 1-3 individuals from 13 of our 14 populations. We obtained a single estimate of genome size for 13-28 additional individuals from across those 14 populations, for a total 333 individuals for which we have at least 1 genome size estimate (Table 1).

### Genome size correction

Preliminary analyses of the genome size data indicated that the date of flow cytometry measurement had a weakly negative, but highly significant, effect on estimated genome size (see Results, Table 2). This trend might reflect variation induced by developmental changes in the plant as it ages or changes in the flow cytometer over time (Doležel and Bartoš 2005; Doležel *et al*. 2007). We calculated a corrected genome size estimate using the coefficient from the best fitting linear model predicting genome size from age of the plant at measurement, in which corrected genome size = genome size estimate - (−0.00031*days between germination and estimate). We used the date-corrected genome size in all subsequent trait analyses.

**Table 2.** Linear model explaining 2C genome size (pg) variation. Significant effects are shown in bold. Effects without significant main or interaction effects (*p* >0.1) were removed from the model (NS).

| Effect type | Effect | Coefficient | df | F-ratio | $p$ |
| --- | --- | --- | --- | --- | --- |
| Fixed | **population age** | **-0.00030** | **1** | **13.763** | **0.0002** |
| Fixed | elevation | -0.00001 | 1 | 3.567 | 0.0598 |
| Fixed | **days until FC<sup>a</sup>** | **-0.00031** | **1** | **13.022** | **0.0004** |
| Fixed | population age*elevation |  |  |  | NS |
| Fixed | population age*days until FC |  |  |  | NS |
| Fixed | elevation*days until FC |  |  |  | NS |
| Fixed | pop age*elevation*days until FC |  |  |  | NS |
|  |  |  |  | R<sup>2</sup> | 0.0784 |
|  |  |  |  | F<sub>(3,329)</sub> | 9.324 |
| | | | | $p$ | <0.0001 |

<sup>a</sup> number of days between germination and date of flow cytometry estimation

### Statistical analyses

The growth and flowering traits that we collected are likely to be correlated as part of plant development, and so we used a principal component analysis (PCA) to identify major axes of variation in these traits using base R (R Core Team, 2021). Coordinates of PC1 and PC2 were then used as response variables in linear models of the effect of genome size on traits. We fit separate linear models to predict PC1 and PC2 from fixed effects of corrected genome size, greenhouse position, days until harvesting, and interactions among all fixed effects. Greenhouse position was treated as a fixed ordinal variable, reflecting the distance from the evaporative cooling system at one end of the greenhouse, due to a temperature gradient of ∼8°C across our experiment. For both analyses, model selection was performed using analyses of variance (ANOVA) to identify if the removal of non-significant effects (*p*>0.1) significantly improved the model.

We used linear models to test our hypotheses that genome size declines toward the leading edge of invasion and at high elevations within the Californian invasion. Corrected genome size (N=333) was modeled as a function of fixed effects of age of source population, elevation, and position in the greenhouse. We used the Jepson Online Herbarium (http://ucjeps.berkeley.edu/) for *C. solstitialis* in California to determine population age as in Braasch *et al*. (2019). We used the earliest occurrence record in the county of each source population, or the record for a neighbouring county if the collection site was closer to an older occurrence record, as its colonization date. We found elevation using Google Earth records from latitude and longitude of collection locations. The analysis was repeated for genome size estimates that had 3 replicates (N=24), using the mean value across replicates.

## Results

Our survey of populations in the Californian invasion identified intraspecific 2C genome size variation of 1.21 fold across individuals, ranging from 1.58-1.85pg (1.47 - 1.77 GB), with a mean of 1.72pg (1.68 GB) across all genome size measurements (Fig. 2). In our reduced dataset of only individuals with repeated estimates, we determined the means across each individual’s estimates and recovered the same range of genome size from 1.58-1.85pg (Supplemental Fig. S1), with a slightly greater mean of 1.74pg (1.70 Gbp). We also found a high degree of within-population variation in genome size, such that most populations included genome sizes spanning the range of the dataset. For example, the coastal Bay area populations MAR and NAP had the greatest intrapopulation range of 1.58-1.78pg (1.55-1.74 GB) and 1.60-1.80pg (1.56-1.76 GB), respectively, and the Central Valley population RB had the narrowest range in genome sizes at 1.66 - 1.76pg (1.62-1.72 GB).

**Figure 2.**
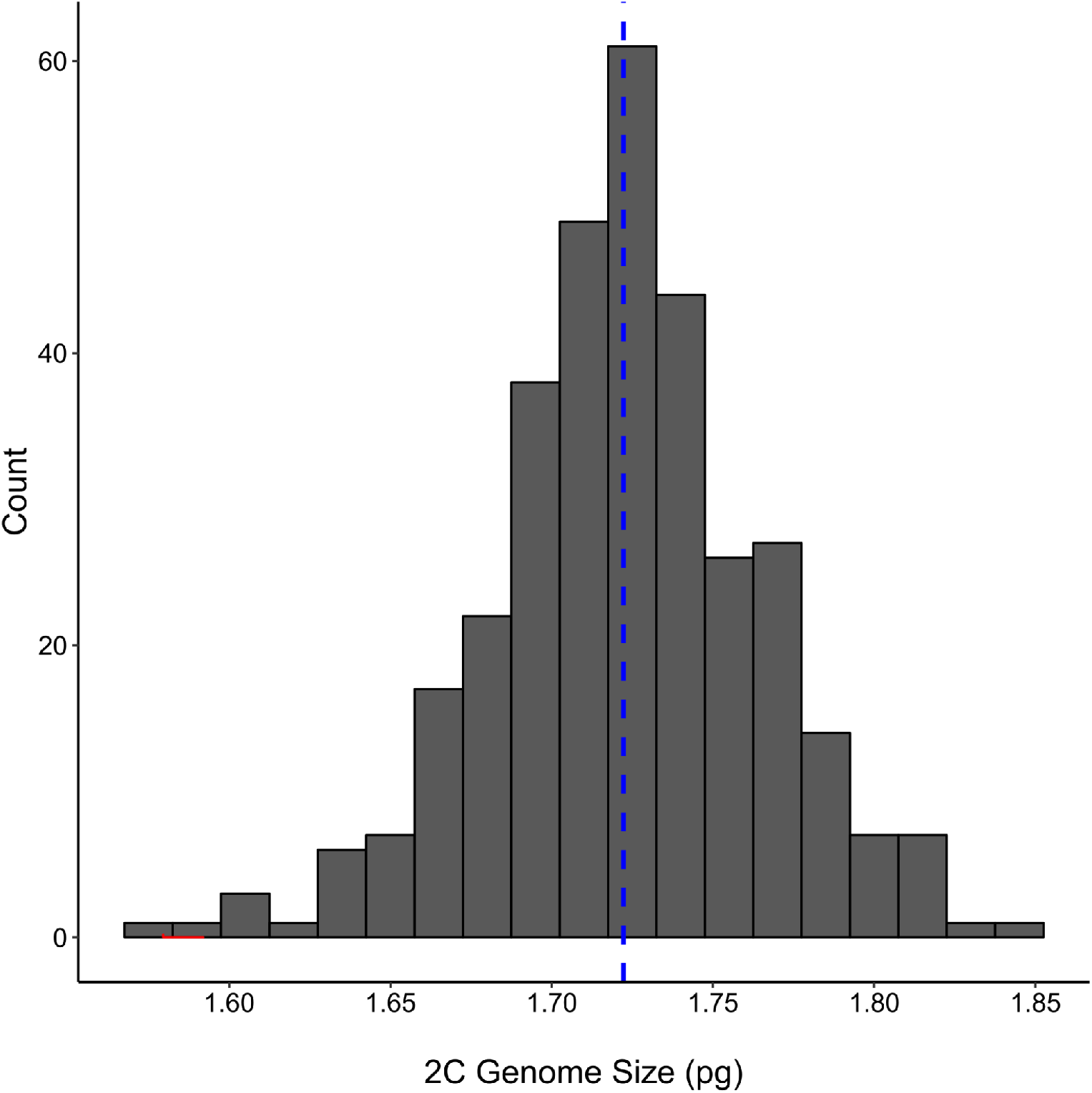
Histogram of 2C genome sizes sampled from the Californian invasion (N=335), with a mean size of 1.72pg (blue dashed line). For individuals with repeat measurements (N=24), only the means of estimated genome sizes are included.

The best-fitting linear model explaining variation in genome size identified significant negative effects of population age, such that more recently founded populations had higher genome sizes, consistent with a scenario of reduced efficacy of selection in recently established populations (Fig 3a, Table 2). The effect of elevation had a marginally significant, negative relationship with genome size, consistent with selection for faster growth and early reproduction at higher elevations, where growing seasons tend to be shorter (Fig 3c, Table 2). We repeated the analysis of genome size with 24 individuals for which we had 3 replicate estimates. We again found a significant negative relationship with population age (Fig 3b); however, there was no significant relationship with elevation in the reduced dataset, and no effect of genome size estimation date.

**Figure 3.**
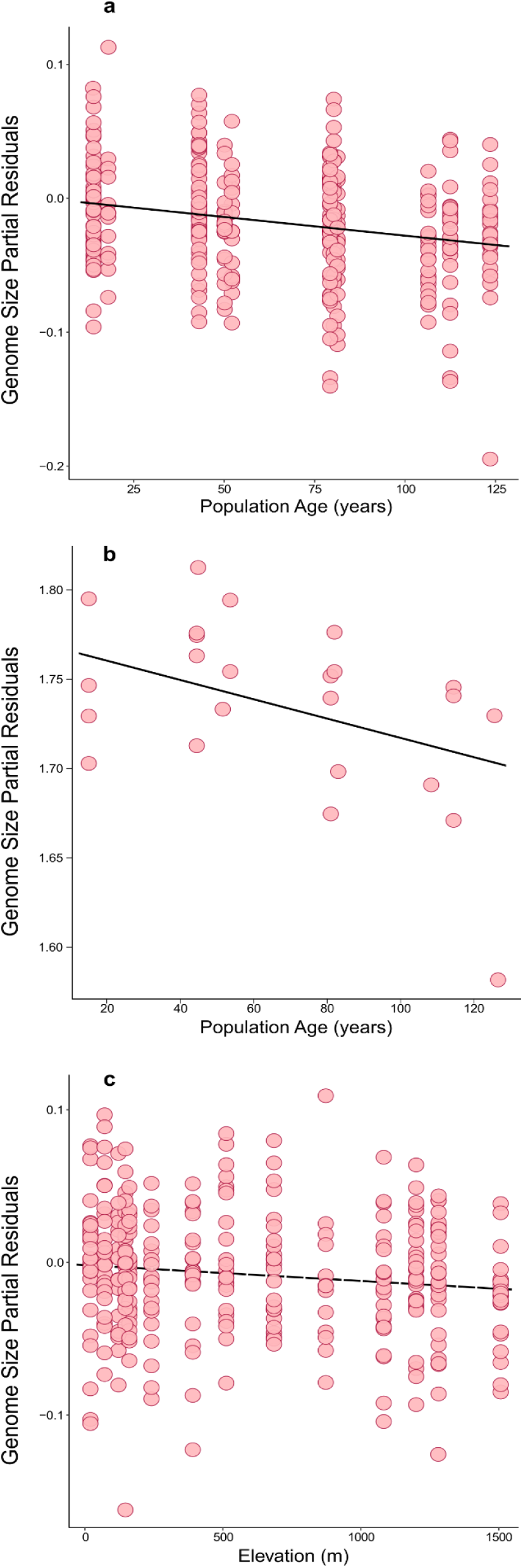
Partial residuals from best-fitting linear models explaining patterns of genome size variation using the full genome size data set of California individuals (N=335) by (a) population age and (c) elevation. (b) Partial residuals by population age using a reduced dataset of only individuals with repeat flow cytometry measurements (N=24). Solid lines indicate significant association (p<0.05), dashed line indicates a marginally significant association (p<0.1).

As noted in the Methods, wee identified a significant negative association between genome size and estimation date, in which flow cytometry measurements later in the growing season yielded significantly smaller estimates of genome size (*β*=-0.00031, *p*<0.001). We used the estimation date coefficient from our best-fitting model explaining genome size variation (Table 2) to apply the following correction:

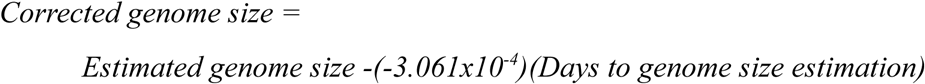

Subsequent models assessing trait variation included this corrected genome size in place of the raw estimates.

Among growth and flowering traits, the first two principal components explained 67.3% of total variance, with PC1 and PC2 accounting for 40.9% and 26.4% variance, respectively (Fig. 4). Principal component analysis indicated that PC1 was strongly associated with growth rates and aboveground dry biomass, while PC2 was associated with bolting time, flowering time, and flower number at harvest (Fig. 4). Higher PC1 values corresponded to faster growth and larger overall size. Higher PC2 values corresponded to longer time to bolting and flowering, and fewer flowers produced overall. PC1 and PC2 coordinate values were used as measures of growth and development metrics for subsequent analyses of trait variation.

**Figure 4.**
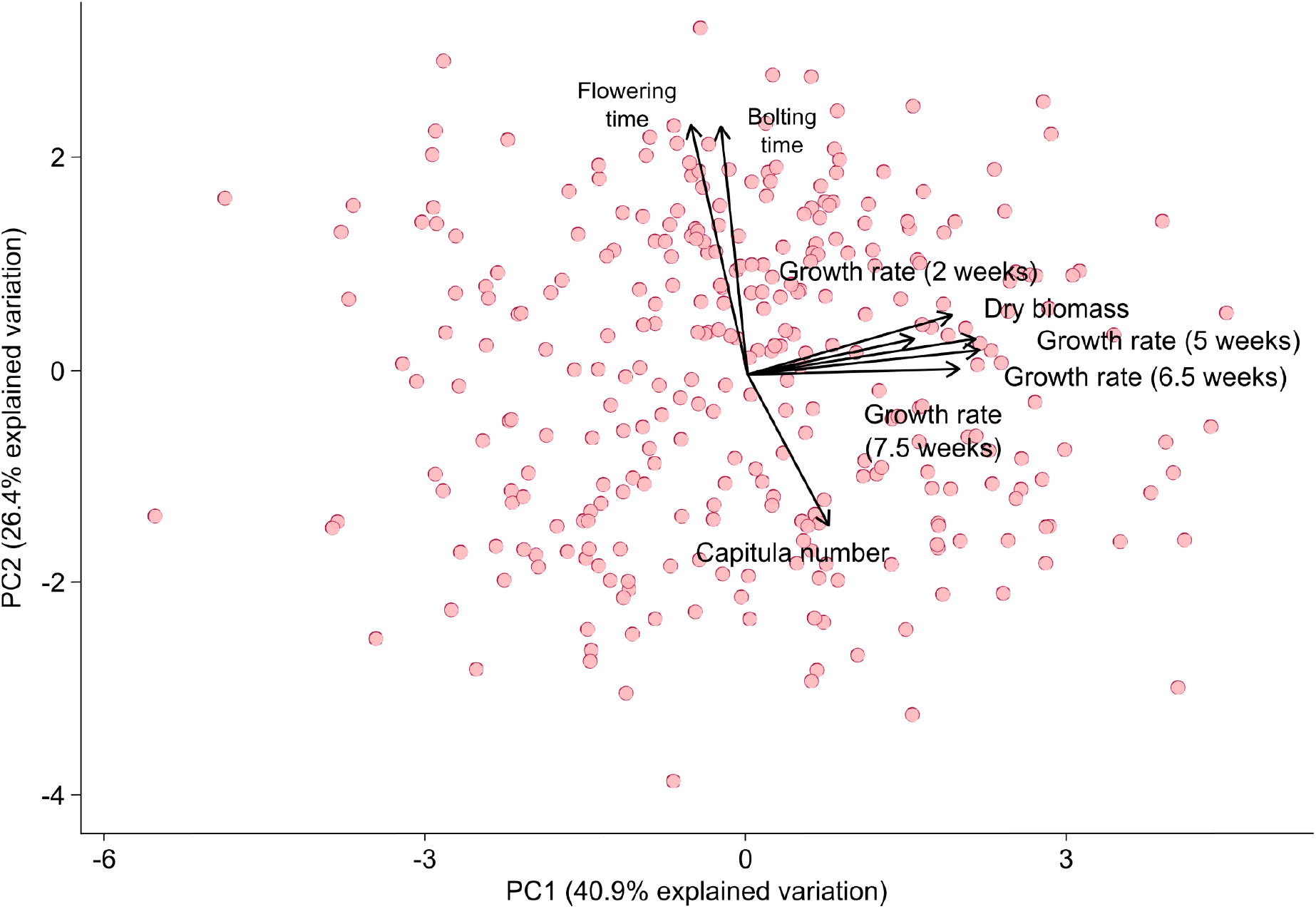
Principal component analysis of traits where each point indicates an individual plant in the common garden (N=391) with PC1 and PC2 explaining 67.3% of all variation. PC1 is associated with variation in growth traits and PC2 is associated with variation in development and reproduction.

In total, we collected both flow cytometry and trait data from 312 individuals. A linear model explaining variation in PC1 scores included non-significant main effects of corrected genome size, with significant interactions between genome size and greenhouse position (Table 3). A partial residual plot of PC1 versus genome size shows a positive slope (Fig 5a), despite a negative coefficient predicted by the linear model on the effect of genome size alone (Table 3). This discrepancy occurred because the effect of genome size on growth-related traits was dependent on where in the greenhouse an individual was grown (Fig. 6). There were no significant effects of genome size on PC1 in positions 1-6, where environmental conditions were warmer, but positions 7 and 8 demonstrated significant associations with genome size where conditions were cooler (SI Table 1). We also found the direction of the relationship between genome size and growth-related traits was more positive in blocks that were cooler, as in the case of position 9 (Fig 6). A linear model explaining variation in PC2 indicated a significant positive effect of corrected genome size (*β*=6.476, *p*=0.001) and negative effects of greenhouse position and its interaction with harvesting time (Table 4, SI Table 2). Genome size was significantly, positively correlated with the timing of reproductive events, wherein higher PC2 axis values indicate a longer time to bolting and flowering, as well as lower capitula production (Fig. 5b), consistent with the prediction that genome replication costs and longer cell cycles may delay reproduction.

**Table 3.** Best-fitting linear model explaining PC1 coordinates (composite of growth rate and biomass traits). Significant effects are in bold. Effects without significant main or interaction effects (*p* >0.1) were removed from the model (NS).

| Effect type | Effect | Coefficient | df | F-ratio | $p$ |
| --- | --- | --- | --- | --- | --- |
| Fixed | genome size<sup>a</sup> | -6.752 | 1 | -0.857 | 0.392 |
| Fixed | **greenhouse position<sup>b</sup>** | **see SI<sup>c</sup>** | **8** | **3.411** | **0.0009** |
| Fixed | **genome size\*greenhouse position** | **see SI<sup>c</sup>** | **8** | **2.008** | **0.0454** |
| Fixed | days to harvest |  |  |  | NS |
| Fixed | genome size*days to harvest |  |  |  | NS |
| Fixed | greenhouse position*days to harvest |  |  |  | NS |
| Fixed | higher order interactions |  |  |  | NS |
| Random | block |  |  |  | NS |
|  |  |  |  | R<sup>2</sup> | 0.126 |
|  |  |  |  | F<sub>(17,294)</sub> | 2.481 |
| | | | | $p$ | 0.001 |

<sup>a</sup> 2C genome size corrected for effect of estimation date
<sup>b</sup> distance to cooling pads in greenhouse, scored as positions 1 (furthest) - 9 (closest)
<sup>c</sup> see Supplementary information (SI) for effect level and interaction coefficients

**Table 4.**
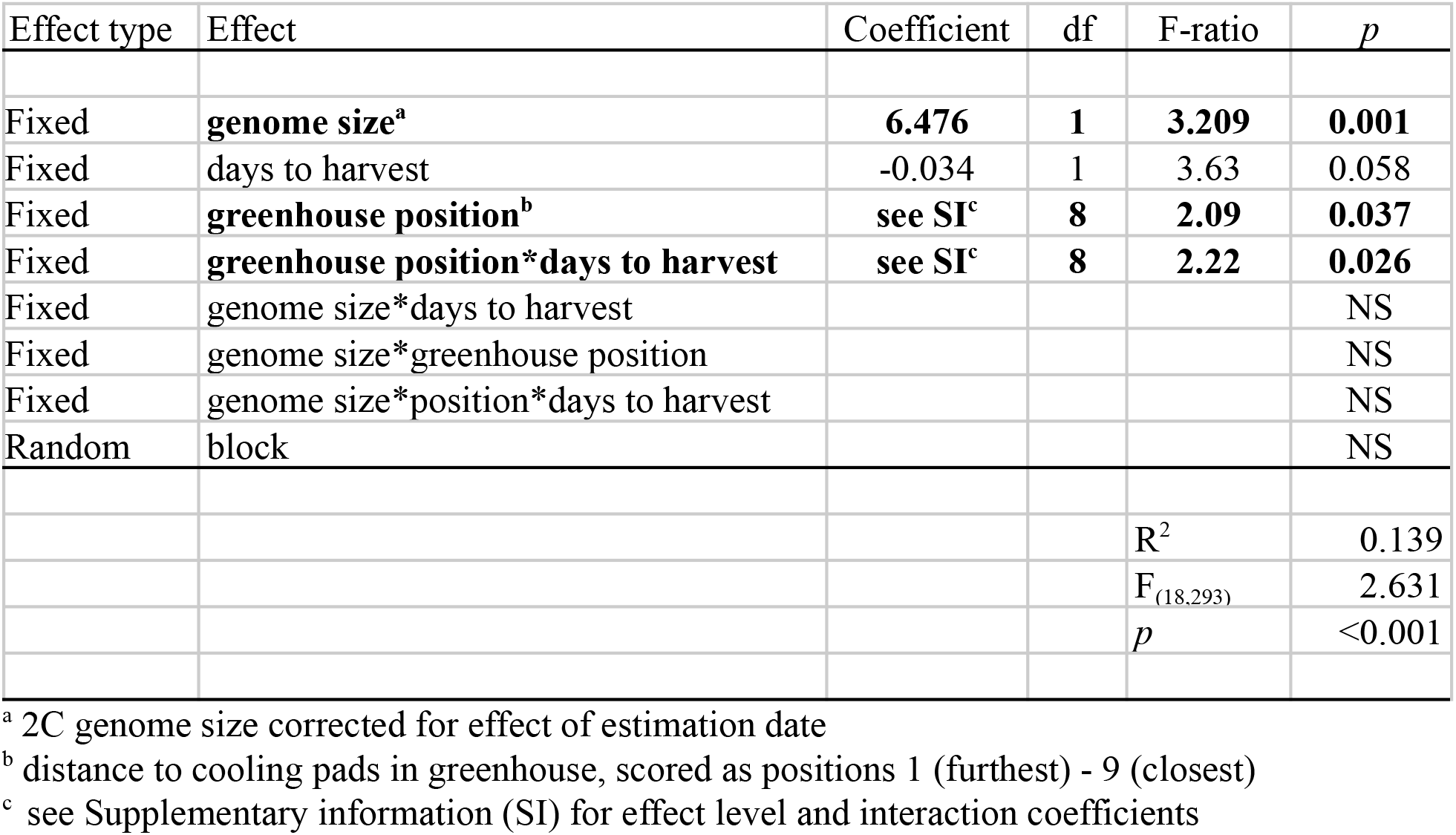
Best-fitting linear model explaining PC2 (composite of development time and flowering traits). Significant effects are shown in bold. Effects without significant main or interaction effects (*p* >0.1) were removed from the model (NS).

| Effect type | Effect | Coefficient | df | F-ratio | $p$ |
| --- | --- | --- | --- | --- | --- |
| Fixed | **genome size<sup>a</sup>** | **6.476** | **1** | **3.209** | **0.001** |
| Fixed | days to harvest | -0.034 | 1 | 3.63 | 0.058 |
| Fixed | **greenhouse position<sup>b</sup>** | **see SI<sup>c</sup>** | **8** | **2.09** | **0.037** |
| Fixed | **greenhouse position\*days to harvest** | **see SI<sup>c</sup>** | **8** | **2.22** | **0.026** |
| Fixed | genome size*days to harvest |  |  |  | NS |
| Fixed | genome size*greenhouse position |  |  |  | NS |
| Fixed | genome size*position*days to harvest |  |  |  | NS |
| Random | block |  |  |  | NS |
| | | | | $R^2$ | 0.139 |
| | | | | $F_{(18,293)}$ | 2.631 |
| | | | | $p$ | <0.001 |

<sup>a</sup> 2C genome size corrected for effect of estimation date
<sup>b</sup> distance to cooling pads in greenhouse, scored as positions 1 (furthest) - 9 (closest)
<sup>c</sup> see Supplementary information (SI) for effect level and interaction coefficients

**Figure 5.**
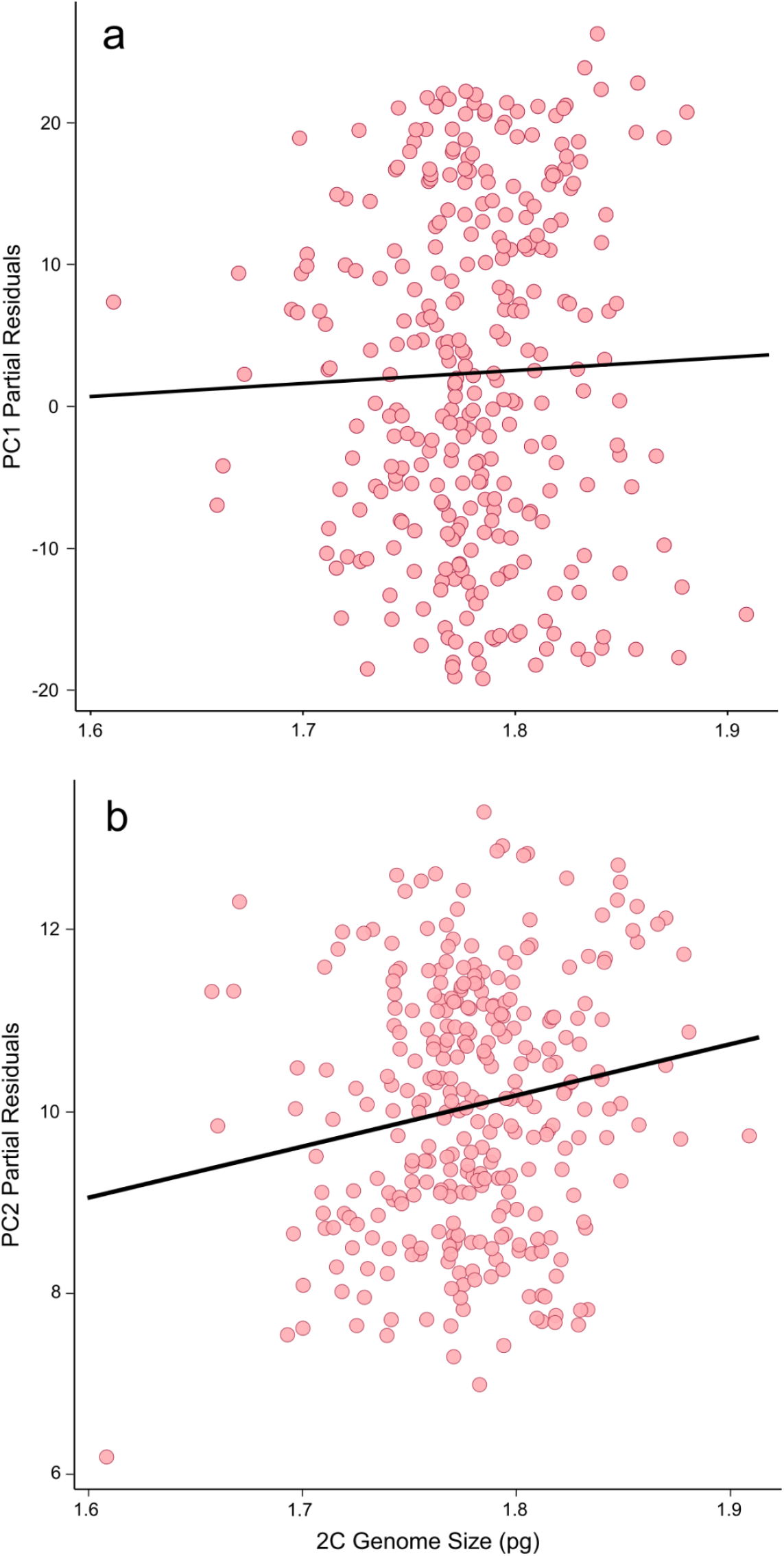
(a) Partial residual plot from best-fitting model explaining variation in PC1 scores due to fixed effect of genome size (NS, *p*=0.39). (b) Partial residual plot explaining variation in PC2 scores by genome size (*p*=0.001)

**Figure 6.**
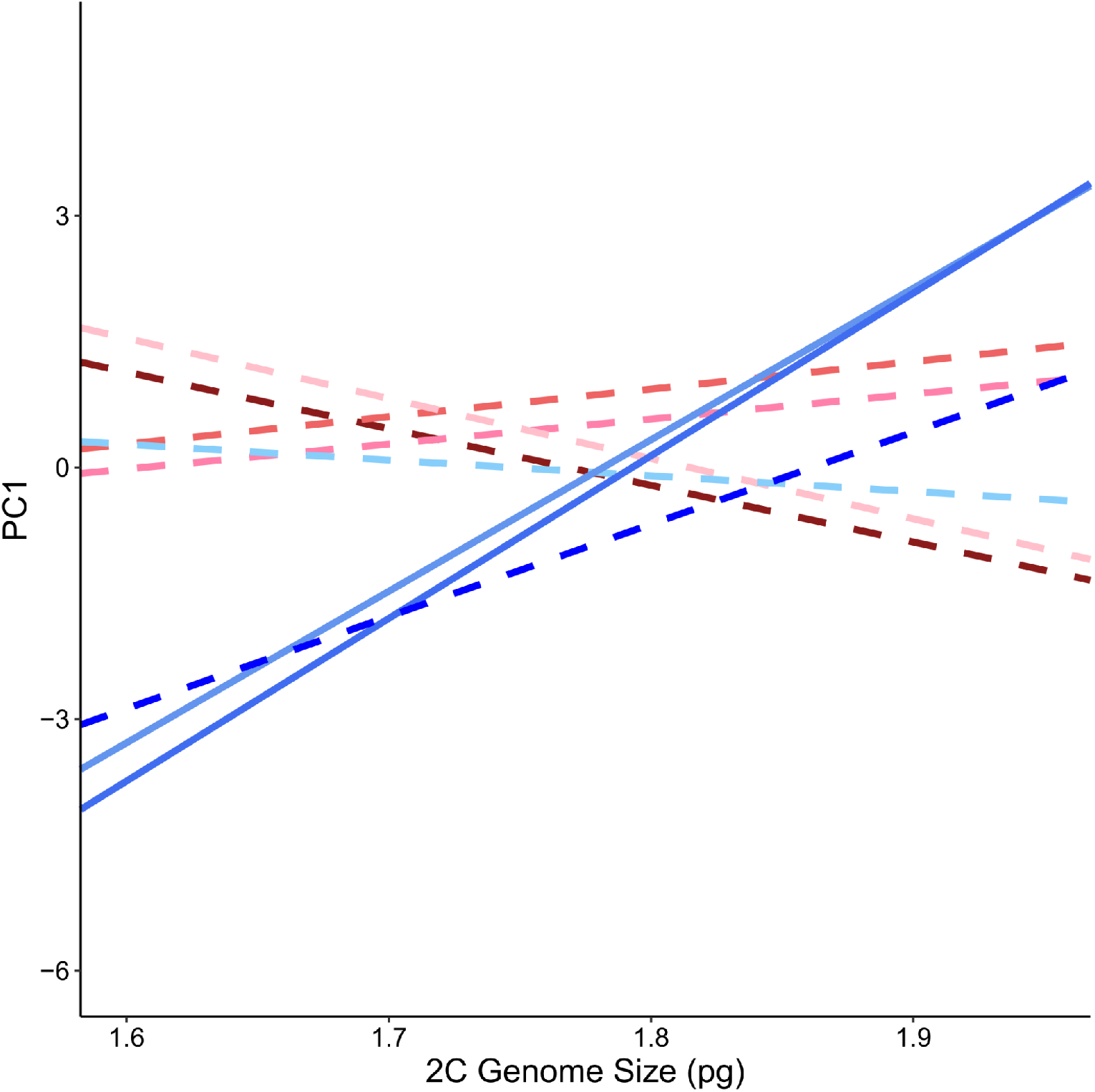
Genome size varies in direction and magnitude of effect on PC1 scores when plotted by greenhouse block. Different colour lines from red to blue indicate greenhouse position with increasing proximity to the cooling pads, where solid lines indicate a significant effect of genome size on PC1 (*p*<0.05). Significant and marginally significant positive correlations were found in positions 7 and 8, and position 9, respectively, where local environments were coolest.

## Discussion

Intraspecific homoploid genome size variation has been demonstrated in several plant systems, but the ecological and evolutionary factors shaping this variation, and its potential contribution of this genetic variation to species invasions, remain unclear. We conducted a survey of genome size and trait variation in YST populations across its invasion in California to test alternative eco-evolutionary hypotheses explaining divergence in genome size during range expansion. The 2C DNA content varied 1.21-fold among all California samples, suggesting widespread and extensive genome size variation across this invasion. Genome size variation was associated with flowering time variation, such that plants with larger genomes reproduced later, with lower lifetime capitula production. Genome sizes increased toward the leading edge of the invasion, but tended to decrease at higher elevations, consistent with genetic drift during range expansion but potentially strong selection for smaller genomes and faster development time at higher elevations.

Surveys of intraspecific genome size variation in other systems have documented both broad-scale variation and conserved genome sizes across populations. Natural populations can produce intraspecific monoploid genome sizes that are almost twice the size or more of the smallest reported variants, for example 1.7-fold in *Brachypodium distachyon* (Oney-Birol and Tabur 2018), and over 2-fold in brown algae *Synura petersenii* (Certnerova and Skaloud 2020). Conversely, genome size can also be a very stable species-level trait with little observed variation, despite widespread population sampling (eg. Lysak *et al*. 2000; Savas *et al*. 2019). As our Californian samples varied 1.21fold, YST demonstrated moderate variation on par with a previous survey in the system (Irimia *et al*. 2017) and with observations in several other systems including *Dactylis glomerata* (Creber *et al*. 1994), *Arachis duranensis* (Temsch and Greilhuber 2001), *Lagenaria siceraria* (Achigan-Dako *et al*. 2008) and *Phragmites australis* (Pysek *et al*. 2018). The variation we observed in YST is also similar to systems that have demonstrated significant associations between genome size and important plant traits (Pysek *et al*. 2018), or with environmental variables (Benor *et al*. 2007; Achigan-Dako *et al*. 2008; Bilinski *et al*. 2018). Our means of triplicate genome size estimates also span this range of variation, and bracket previous single estimates (Bancheva and Greilhuber 2006; Miskella 2014; Carev *et al*. 2017), indicating that measurement errors are unlikely to explain the 1.21-fold variation observed across our samples (Greilhuber 2005).

Notably, we detected an influence of measurement date on genome size estimates, in which later estimates tended to yield smaller genome sizes. Flow cytometry measurements that underestimate nuclear size can occur when DNA staining is inhibited (Doležel *et al*. 2007). Genome size measurements were taken over five months, during which the plants progressed from rosette formation in early development, through early senescence. Phenolic concentrations in the cytosol can vary among tissue types and across developmental stages (Witzell *et al*. 2003; Wam *et al*. 2017), some of which are known to inhibit the staining of DNA (Doležel *et al*. 2007). Fortunately, our relatively large sample size allowed us to identify this artifact and most importantly, use our linear model explaining genome size variation to quantify the effect and remove it from subsequent analyses of trait variation.

We found that this broad genome size variation was significantly associated with variation in reproductive traits, wherein larger genomes were associated with later flowering and lower lifetime capitula production. These results join a growing body of evidence that points to potential developmental costs of maintaining large genomes. Across a diversity of species, genome size has been correlated with a suite of characteristics, including plant size (Munzbergova 2009; Carta and Peruzzi 2016), growth rate (Fridley and Craddock 2015), pollen tube growth (Reese and William 2019), seed size (Beaulieu *et al* 2007), flowering time (eg. Benor *et al*. 2011; Jian *et al*. 2017), and rates of photosynthesis (Roddy *et al*. 2020). Comparative phylogenetic approaches to identifying the effects of genome size on traits have found that genome size can be a strong predictor of seed mass (Knight and Beaulieu 2008), which may mediate the apparent effect of genome size on other important traits (Grotkopp *et al*. 2004).

Importantly, the magnitude and direction of genome size effects on growth-related traits in our study were dependent on local environmental conditions. Our linear model demonstrated a significant effect of the interaction between greenhouse position and genome size on growth, in which genome size had a significant positive association with growth in blocks that were in cooler locations. The nature of this interaction was predominantly driven by individuals with small genomes that grew more slowly and produced less biomass overall under cooler growing conditions. A similar interaction effect between genome size and variation in greenhouse conditions on aboveground biomass has also been identified in *Phragmites australis* (Meyerson *et al*. 2020). These interactions suggest that the effect of selection on genome size, and the subsequent relationship between genome size and the traits it influences, will vary across environments. For example, field surveys of an experimental grass community in the UK found that species with large genomes were underrepresented in nutrient-poor conditions with lower productivity, but successfully competed with small genome species and produced more biomass in N+P supplemented plots (Guignard *et al*. 2016). Principal component analysis of traits in our study identified that major axes associated with growth and reproduction varied independently from each other, but genome size influenced variation across both axes. These relationships suggest that effects of genome size on overall plant performance and fitness are complex, and can manifest across multiple independent traits and be environmentally-dependent. A critical consideration for additional experiments is that the apparent scale of genome size effects on trait variation may depend on the environments under which they are measured. Variation in the strength of these effects could then be difficult to quantify without explicit testing under varying environments.

We predicted that selection would favor smaller genome sizes during range expansion in habitats that favor faster development times. There is increasing evidence of correlations between genome size and a variety of environmental conditions, such as elevation (Achigan-Dako *et al*. 2008; Benor *et al*. 2011; Diez *et al*. 2013; Carta and Peruzzi 2015; Bilinski *et al*. 2018), temperature seasonality (Knight and Ackerly 2002; Paule *et al*. 2018; Qiu *et al*. 2019), atmospheric carbon (Franks *et al*. 2012), precipitation (Knight and Ackerly 2002; Carta and Peruzzi 2015), heavy metal pollution (Vidic *et al*. 2009; Temsch *et al*. 2010) and soil nitrogen (Kang *et al*. 2015; Guignard *et al*. 2016). We found marginally significant evidence for small genome sizes at high elevations in our full dataset. However, the marginally significant relationship with elevation disappeared in the reduced dataset, as did the effect of date of genome size estimation. These replicates were taken early in YST development, and so variation in age-specific cytosolic compound concentrations likely exerted less influence on estimation precision. The relationship between genome size and elevation is thought to be mediated by selection for rapid onset of flowering due to short growing seasons at high elevations, as appears to be the case in maize (Bilinski *et al*. 2018). We do observe a significant negative correlation between flowering time and elevation, such that strong direct selection on flowering time might explain weaker patterns of genome size differentiation, given that genome size differences explain only a portion of flowering time trait variation.

In contrast, we did not find evidence that genome size declines during range expansion, as we predicted if selection favored fast development times in invaders. Rather, we found that genome sizes increased toward the leading edge of the expansion, a pattern more consistent with founder effects and genetic drift contributing to the accumulation of larger genomes in populations at the leading edge. Repeated analysis of genome size with individuals for which we had 3 replicate estimates confirmed a significant negative relationship with population age. Previous work in this system demonstrated that more recently founded populations have lower effective population sizes, consistent with stronger genetic drift (Braasch *et al*. 2019). Recent theoretical work on the formation of range limits predicts that mutational load resulting from accumulation of deleterious variants during successive founding events can reduce population expansion across an environmental gradient, especially when mutations are not severely deleterious (Henry *et al*. 2015). A survey of genome sizes in European beech recovered similar findings, in which genome sizes increased towards range margins (Paule *et al*. 2018). This has particular importance for our understanding of the evolution of range limits in introduced species, as relatively strong genetic drift could allow mildly deleterious increases in genome size to accumulate and slow expansion at the leading edge. Geographic patterns of mutation accumulation in *Arabidopsis lyrata* support this prediction, in which signatures of mutational load increased towards range edges, with associated effects on individual plant performance and population growth (Willi *et al*. 2018). Therefore, small effective population sizes could explain both how larger genomes in YST persist at the edge of expansion in spite of potential reproductive costs, and why a putatively adaptive relationship with elevation is weak.

Successful management of species invasions and prevention of future spread depends on understanding the factors that shape the dynamics of range expansion. There is a long history in invasion biology of seeking traits that are associated with invaders, but we have only recently begun to investigate structural genomic variation (Kuester *et al*. 2014; Schrader *et al*. 2014; Stapley *et al*. 2015; Gourbet *et al*. 2017; Bilinski *et al*. 2018; Pysek *et al*. 2018; Baduel *et al*. 2019). Our data contribute to increasing evidence that intraspecific genome size variants are common and can generate genetic and phenotypic variation on very short timescales, which may be important to establishment and range edge formation during invasions.

## Acknowledgements

We thank Abreeza Zegeer for advice on greenhouse experimental set up and management. We thank Paula Campbell and John Finch from the University of Arizona Cytometry Core Facility, and Anthony Baniaga for their assistance preparing flow cytometry protocols for genome size estimation. We would also like to thank Joseph Braasch for supplying dates of population establishment and helpful feedback. We are most grateful to Rhiannon Bauer, Alexus Cazares, Shelby Dolgaard and Alex Skomro for their indispensable help with collecting common garden data. Funding was provided by USDA grant #2015-67013-23000 and NSF grant #1750280 to KMD.

## Author Contributions

FAC, SRW and KMD designed the research. FAC, SRW, JWW and MZ collected common garden and flow cytometry data. FAC and KMD conducted data analysis and interpretation. FAC and KMD wrote the manuscript.

## Supplementary Information

**Figure S1.**
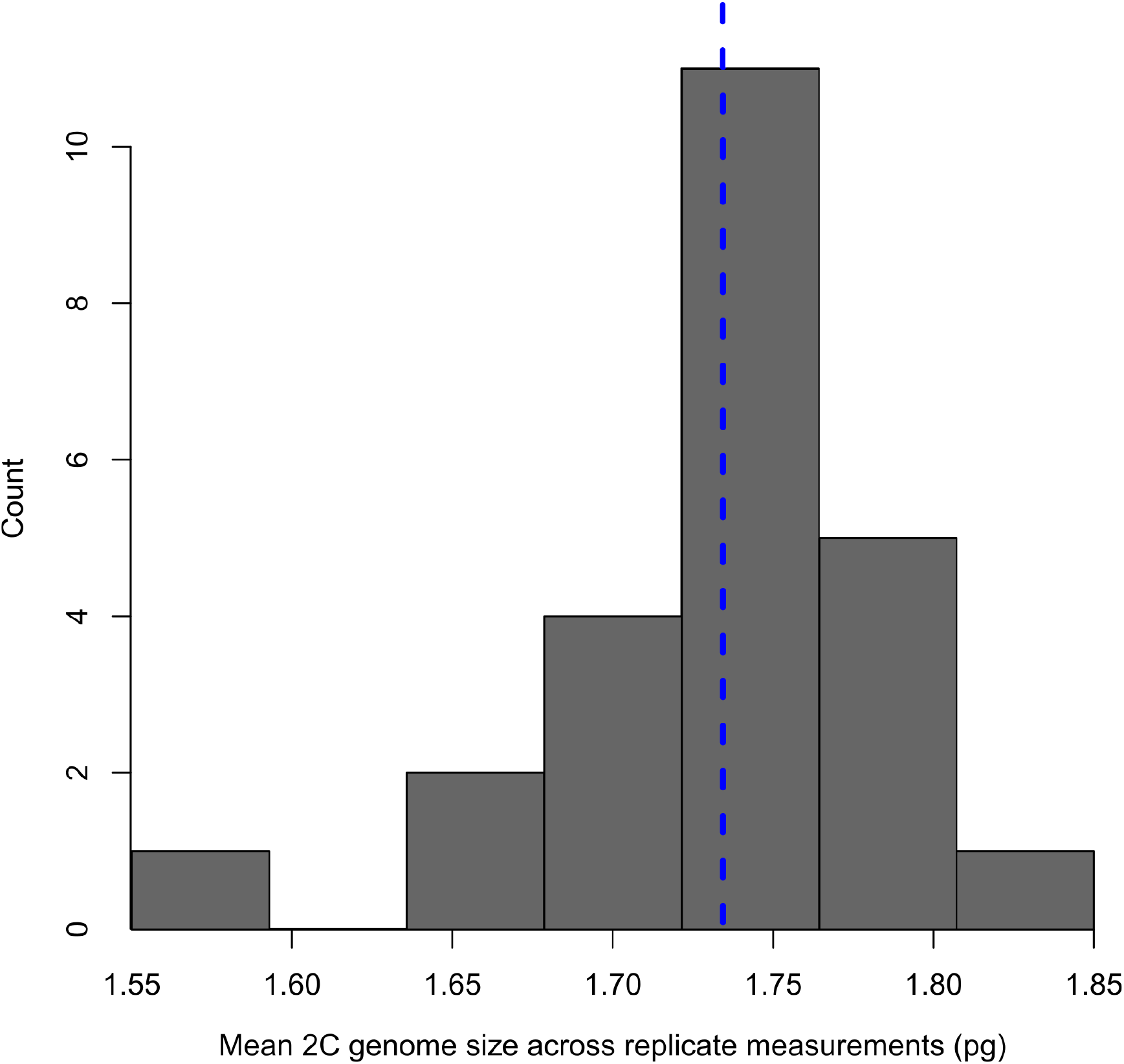
Distribution of within-individual mean genome sizes, including only those that had 3 replicate measurements (N=24). Dashed line is the mean genome size across individuals (mean=1.734pg).

**Table S1.**
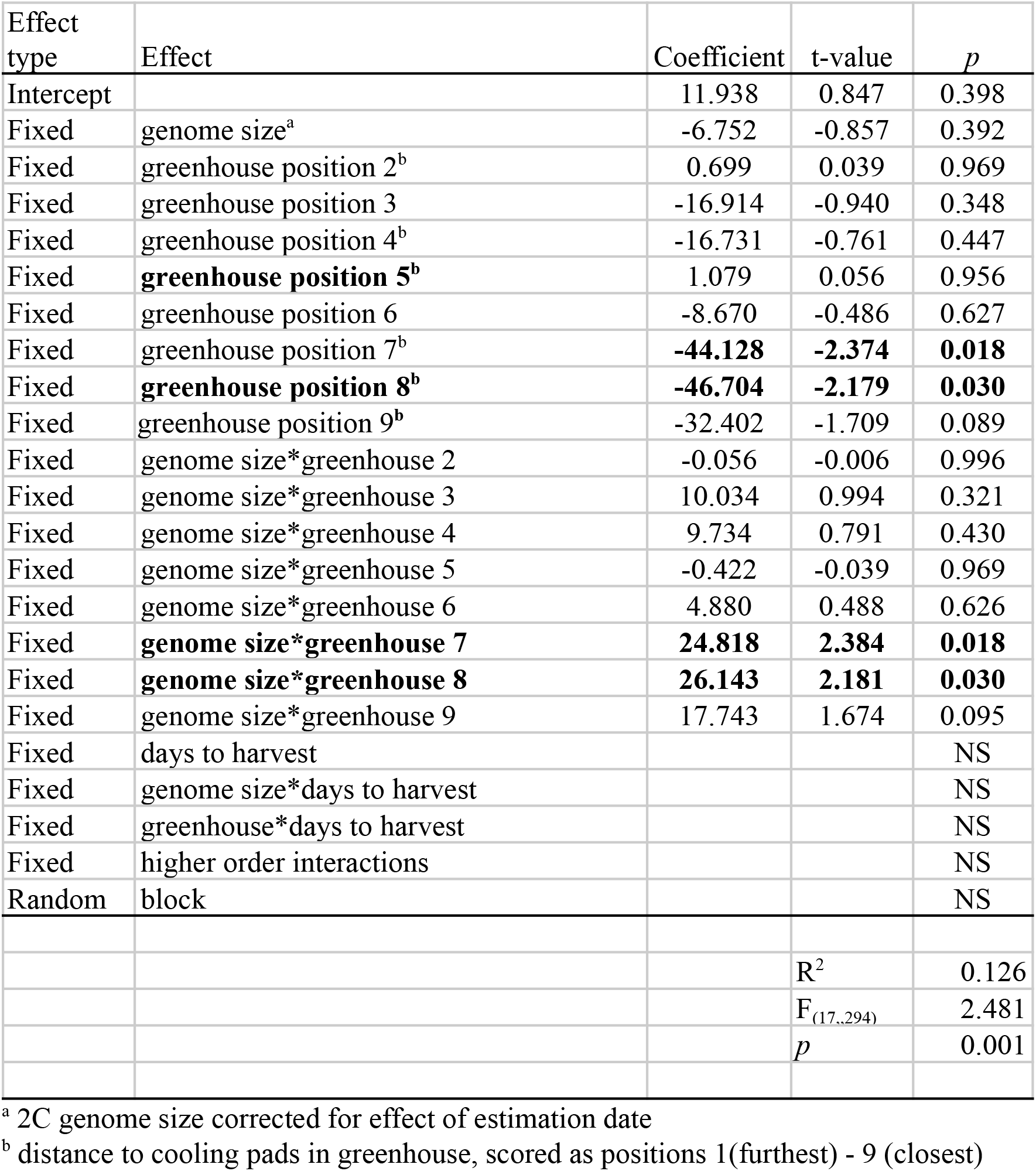
Coefficient details for full best-fitting linear model explaining PC1 (growth-related traits). Significant effects are bold, and effects without significant main or interaction effects were removed.

**Table S2.**
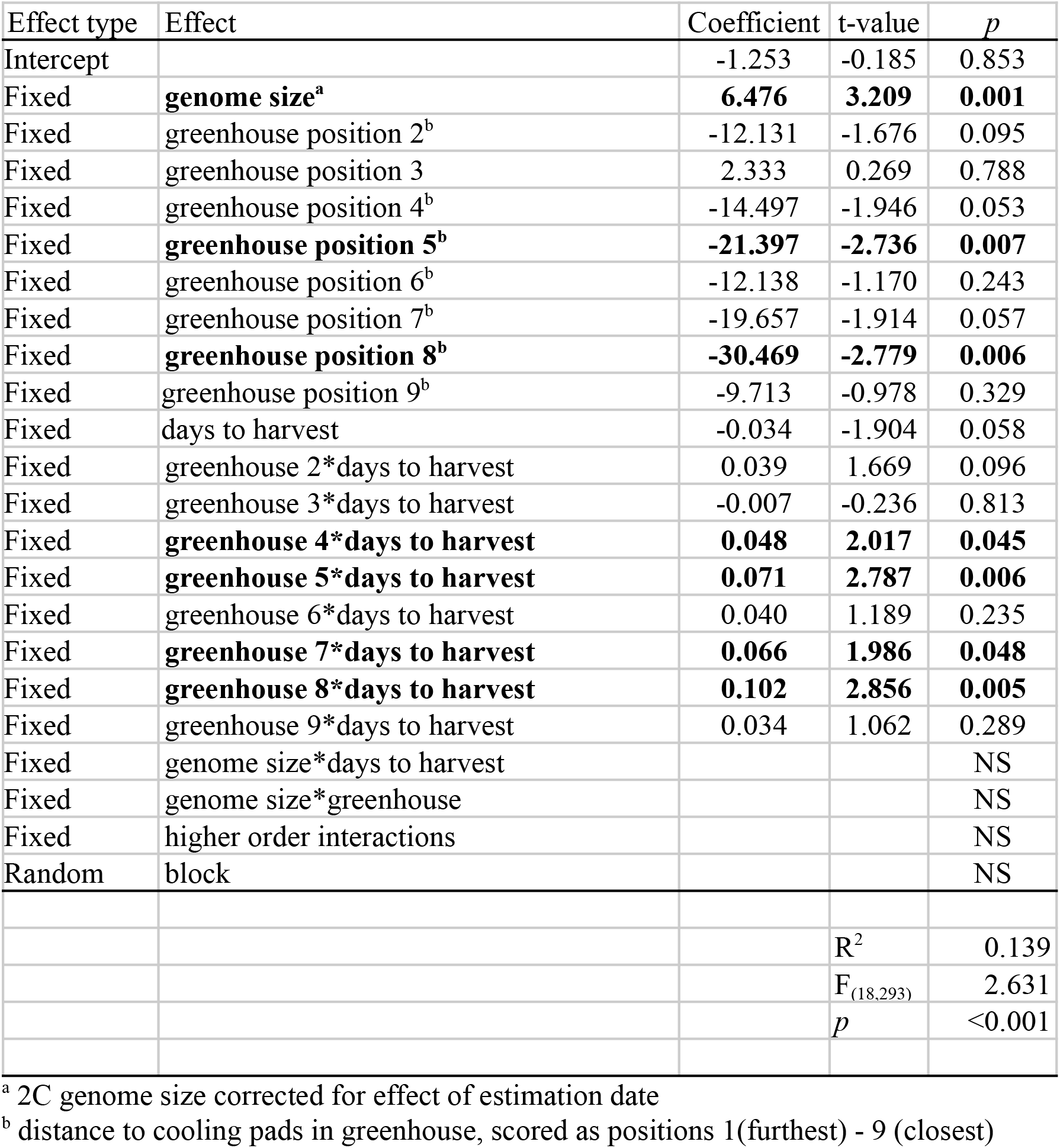
Coefficient details for full best-fitting linear model explaining PC2 (development-related traits). Significant effects are shown in bold, and effects without significant main or interaction effects were removed.

