## Supplementary Information for "Genome size variation and evolution during invasive range expansion in an introduced plant"

**Supplemental Tables and Figures**

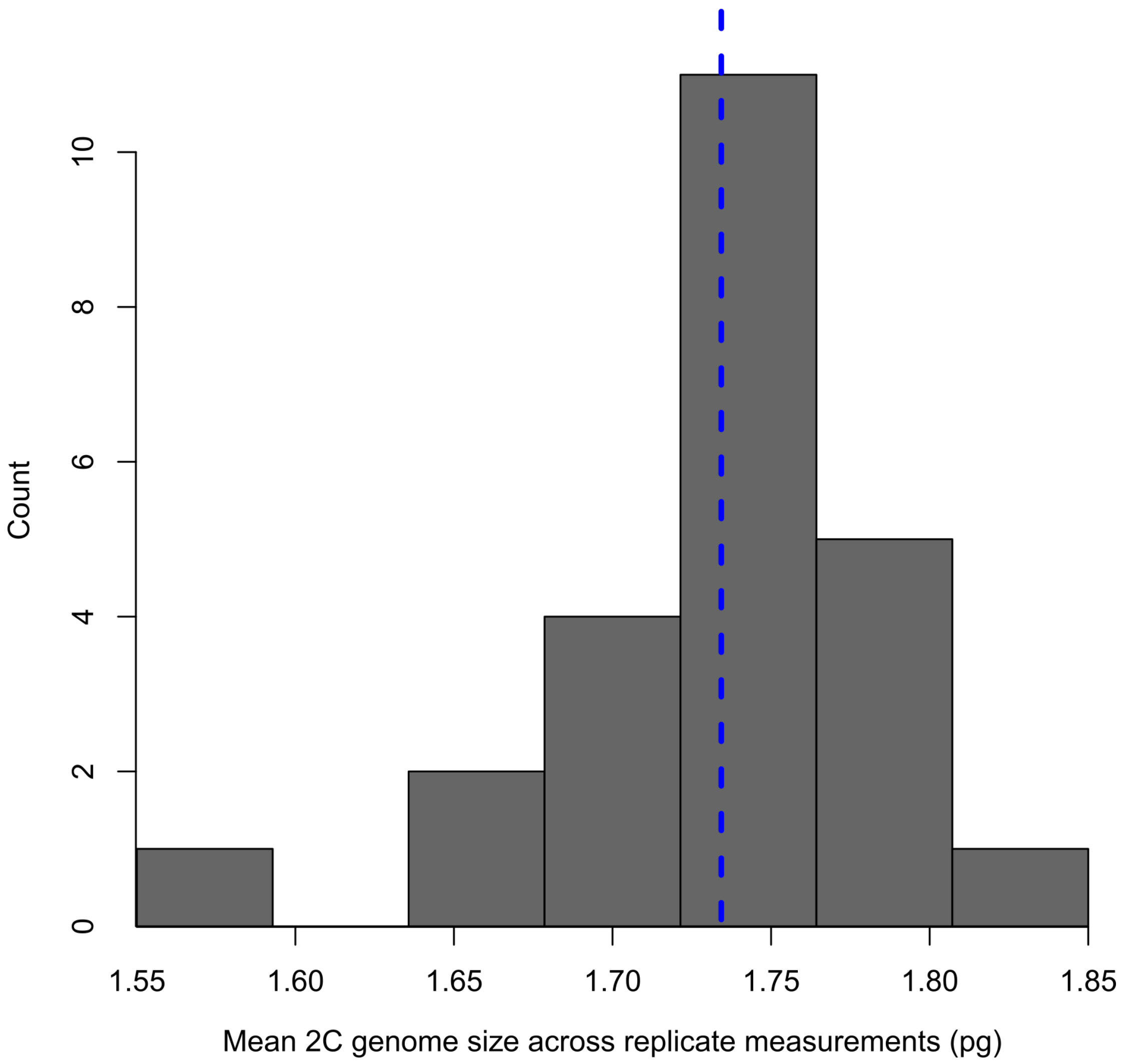

**Figure S1.** Distribution of within-individual mean genome sizes, including only those that had 3 replicate measurements (N=24). Dashed line is the mean genome size across individuals (mean=1.734pg).

**Table S1.** Coefficient details for full best-fitting linear model explaining PC1 (growth-related traits). Significant effects are bold, and effects without significant main or interaction effects were removed.

| Effect type | Effect | Coefficient | t-value | *p* |
|---|---|---|---|---|
| Intercept | | 11.938 | 0.847 | 0.398 |
| Fixed | genome size[a] | -6.752 | -0.857 | 0.392 |
| Fixed | greenhouse position 2[b] | 0.699 | 0.039 | 0.969 |
| Fixed | greenhouse position 3 | -16.914 | -0.940 | 0.348 |
| Fixed | greenhouse position 4[b] | -16.731 | -0.761 | 0.447 |
| Fixed | **greenhouse position 5[b]** | 1.079 | 0.056 | 0.956 |
| Fixed | greenhouse position 6 | -8.670 | -0.486 | 0.627 |
| Fixed | greenhouse position 7[b] | **-44.128** | **-2.374** | **0.018** |
| Fixed | **greenhouse position 8[b]** | **-46.704** | **-2.179** | **0.030** |
| Fixed | greenhouse position 9[b] | -32.402 | -1.709 | 0.089 |
| Fixed | genome size*greenhouse 2 | -0.056 | -0.006 | 0.996 |
| Fixed | genome size*greenhouse 3 | 10.034 | 0.994 | 0.321 |
| Fixed | genome size*greenhouse 4 | 9.734 | 0.791 | 0.430 |
| Fixed | genome size*greenhouse 5 | -0.422 | -0.039 | 0.969 |
| Fixed | genome size*greenhouse 6 | 4.880 | 0.488 | 0.626 |
| Fixed | **genome size*greenhouse 7** | **24.818** | **2.384** | **0.018** |
| Fixed | **genome size*greenhouse 8** | **26.143** | **2.181** | **0.030** |
| Fixed | genome size*greenhouse 9 | 17.743 | 1.674 | 0.095 |
| Fixed | days to harvest | | | NS |
| Fixed | genome size*days to harvest | | | NS |
| Fixed | greenhouse*days to harvest | | | NS |
| Fixed | higher order interactions | | | NS |
| Random | block | | | NS |
| | | | $R^2$ | 0.126 |
| | | | $F_{(17,,294)}$ | 2.481 |
| | | | *p* | 0.001 |

[a] 2C genome size corrected for effect of estimation date
[b] distance to cooling pads in greenhouse, scored as positions 1(furthest) - 9 (closest)

**Table S2.** Coefficient details for full best-fitting linear model explaining PC2 (development-related traits). Significant effects are shown in bold, and effects without significant main or interaction effects were removed.

| Effect type | Effect | Coefficient | t-value | *p* |
|---|---|---|---|---|
| Intercept | | -1.253 | -0.185 | 0.853 |
| Fixed | **genome size[a]** | **6.476** | **3.209** | **0.001** |
| Fixed | greenhouse position 2[b] | -12.131 | -1.676 | 0.095 |
| Fixed | greenhouse position 3 | 2.333 | 0.269 | 0.788 |
| Fixed | greenhouse position 4[b] | -14.497 | -1.946 | 0.053 |
| Fixed | **greenhouse position 5[b]** | **-21.397** | **-2.736** | **0.007** |
| Fixed | greenhouse position 6[b] | -12.138 | -1.170 | 0.243 |
| Fixed | greenhouse position 7[b] | -19.657 | -1.914 | 0.057 |
| Fixed | **greenhouse position 8[b]** | **-30.469** | **-2.779** | **0.006** |
| Fixed | greenhouse position 9[b] | -9.713 | -0.978 | 0.329 |
| Fixed | days to harvest | -0.034 | -1.904 | 0.058 |
| Fixed | greenhouse 2*days to harvest | 0.039 | 1.669 | 0.096 |
| Fixed | greenhouse 3*days to harvest | -0.007 | -0.236 | 0.813 |
| Fixed | **greenhouse 4*days to harvest** | **0.048** | **2.017** | **0.045** |
| Fixed | **greenhouse 5*days to harvest** | **0.071** | **2.787** | **0.006** |
| Fixed | greenhouse 6*days to harvest | 0.040 | 1.189 | 0.235 |
| Fixed | **greenhouse 7*days to harvest** | **0.066** | **1.986** | **0.048** |
| Fixed | **greenhouse 8*days to harvest** | **0.102** | **2.856** | **0.005** |
| Fixed | greenhouse 9*days to harvest | 0.034 | 1.062 | 0.289 |
| Fixed | genome size*days to harvest | | | NS |
| Fixed | genome size*greenhouse | | | NS |
| Fixed | higher order interactions | | | NS |
| Random | block | | | NS |
| | | | $R^2$ | 0.139 |
| | | | $F_{(18,293)}$ | 2.631 |
| | | | *p* | <0.001 |

[a] 2C genome size corrected for effect of estimation date
[b] distance to cooling pads in greenhouse, scored as positions 1(furthest) - 9 (closest)
